# Deciphering the variation in cuticular hydrocarbon profiles of six European honey bee subspecies

**DOI:** 10.1101/2024.07.05.601031

**Authors:** Daniel Sebastián Rodríguez-León, Aleksandar Uzunov, Cecilia Costa, Dylan Elen, Leonidas Charistos, Thomas Galea, Martin Gabel, Ricarda Scheiner, María Alice Pinto, Thomas Schmitt

**Affiliations:** University of Würzburg, Biocenter, Department of Animal Ecology and Tropical Biology, Am Hubland, 97074 Würzburg, Germany; Ss. Cyril and Methodius University in Skopje, Faculty for Agricultural Science and Food, 1000 Skopje, Republic of Macedonia; State Key Laboratory of Resource Insects, Institute of Apicultural Research, Chinese Academy of Agricultural Sciences, Beijing, China; CREA Research Centre for Agriculture and Environment, Via di Corticella 133,40128 Bologna, Italy; Bangor University, School of Natural Sciences, Department of Molecular Ecology & Evolution, Bangor LL57 2DG, United Kingdom; ZwarteBij.org vzw, Taskforce Research, 9890 Gavere, Belgium; Hellenic Agricultural Organization DIMITRA, Institute of Animal Science, Department of Apiculture, 63200 Nea Moudania, Greece; Breeds of Origin, Ħaż – Żebbuġ, Malta; LLH Bee Institute Kirchhain, Erlenstraße 9, 35274 Kirchhain, Germany; University of Würzburg, Biocenter, Department of Behavioral Physiology and Sociobiology, Am Hubland, 97074 Würzburg, Germany; Centro de Investigação de Montanha (CIMO), Instituto Politécnico de Bragança, Campus de Santa Apolónia, 5300-253 Bragança, Portugal; Laboratório Associado para a Sustentabilidade e Tecnologia em Regiões de Montanha (SusTEC), Instituto Politécnico de Bragança, Campus de Santa Apolónia, 5300-253 Bragança, Portugal

**Author notes:** Corresponding author (DS Rodríguez-León).

**Keywords:** Social insects, honey bees, ecology, chemical communication, climate adaptation

## Abstract

The Western honey bee (*Apis mellifera*) subspecies exhibit local adaptive traits that evolved in response to the different environments that characterize their native distribution ranges. An important trait is the cuticular hydrocarbon (CHC) profile, which helps preventing desiccation and mediating communication. We compared the CHC profiles of six European subspecies (*A. m. mellifera, A. m. carnica, A. m. ligustica, A. m. macedonica, A. m. iberiensis*, and *A. m. ruttneri*) and investigated potential factors shaping their composition. We did not find evidence of adaptation of the CHC profiles of the subspecies to the climatic conditions in their distribution range. Subspecies-specific differences in CHC composition might be explained by phylogenetic constraints or genetic drift. The CHC profiles of foragers were more subspecies-specific than those of nurse bees, while the latter showed more variation in their CHC profiles, likely due to the lower desiccation stress exerted by the controlled environment inside the hive. The strongest profile differences appeared between nurse bees and foragers among all subspecies, suggesting an adaptation to social task and a role in communication. Foragers also showed an increase in the relative amount of alkanes in their profiles compared to nurses, indicating adaptation to climatic conditions.

## Introduction

Subspecies are populations of organisms that show signs of local adaptation and can be discriminated by morphological, behavioral, and molecular characteristics [1]. The Western honey bee (*Apis mellifera*) is an excellent model to investigate adaptation to local climate because it has evolved into 31 subspecies [2–6]. Although *A. mellifera* is currently distributed worldwide due to anthropogenic dispersion [2, 7–9], its native range comprises Africa and large parts of Eurasia [2, 7, 8, 10, 11]. Throughout this environmentally diverse range, *A. mellifera* expresses a remarkable genetic and phenotypic variation, which was earlier grouped into four evolutionary lineages based on morphological traits [2] and has recently expanded into seven, based on whole genome data [2, 6, 8, 10, 12–14]. Europe harbors an important component of the *A. mellifera* diversity, which is represented by 10 subspecies. This diversity clusters into three divergent morphological lineages, including the Northern and Western European lineage (M), the Central Mediterranean and Southeastern European lineage (C), and the African lineage (A) [2, 4]. Lineage M extends from Southern Iberia, where the summers are very warm and dry, to southern Scandinavia, where the winters are very long and cold. Lineage C comprises the Apennine and Balkan Peninsulas, where the climate ranges from continental (in the north and east) to Mediterranean (in the coastal areas). Finally, lineage A occupies the entire African continent but has historically expanded into Europe, where it is present in Malta and Sicily but also in southeastern Iberia, where the mitochondrial, but not the nuclear DNA, is mostly of African ancestry [2, 15–17].

In this wide geographical area, *A. mellifera* evolved local adaptive traits to thrive across diverse climatic selective pressures [2, 7]. An example of such a trait is the lower thermal tolerance of the European C-lineage *A. m. ligustica* and *A. m. carnica* compared to the Oriental lineage *A. m. jemenitica*, which thrives in the extremely arid habitats of the Arabian Peninsula [18, 19]. Differences in climatic adaptations of honey bee subspecies are particularly important under the current climate change scenario, because they can affect the distribution range and/or drive new competitive relationships between honey bee subspecies or wild bee species [20].

Cuticular hydrocarbons (CHCs) represent an important adaptive trait to climate because they form a semiliquid coating preventing desiccation. In addition, they offer a physical barrier against microorganisms, and mediate intra- and interspecific communication [21–28]. The desiccation barrier of the CHCs depends on the aggregation of the molecules, which, in turn, is determined by the composition of the CHC layer. Tight aggregations are formed by long chain-length hydrocarbons and saturated, non-methyl-branched hydrocarbons (alkanes). The waterproof barrier induced by those hydrocarbons is more efficient than that of shorter chain-length, unsaturated, or methyl-branched hydrocarbons. The latter, however, are known to act as recognition cues for sex, age, nestmate, cast, and task performance [25, 29–40]. The trade-off between water-loss prevention and communication necessities might therefore constrain the variation in CHC compositional profiles [23, 29, 41, 42].

The CHC profiles of honey bee workers display stereotypic qualitative and quantitative variations correlating with age and task performance [31, 37, 43]. These task specific profiles are assumedly an adaptive response of worker bees to contrasting environments (i.e., temperature and humidity).

Workers performing tasks inside the hive (brood care) are naturally exposed to a more controlled environment compared to foraging honey bees, which leave the hive to collect resources and are exposed to a harsher environment. Not surprisingly, foragers have a more waterproof CHC profile than nurse bees [31, 39]. Considering that honey bees can discriminate between different CHC profiles, those differences might serve as recognition cues for task performance among worker bees [31, 43, 44]. Whether the CHC profile compositions exhibited by different *A. mellifera* subspecies is subspecies-specific and adapted to different climatic conditions is an open question.

In the present study, we analyzed the CHC profiles of nurse bees and foragers of six European *A. mellifera* subspecies (*A. m. mellifera* and *A. m. iberiensis* of M-lineage ancestry, *A. m. carnica, A. m. ligustica*, and *A. m. macedonica* of C-lineage ancestry, and *A. m. ruttneri* of A-lineage ancestry) raised under the same environmental conditions. We chose a common-garden experiment to exclude the impact of the local environment on the expression of the potential subspecies-specific CHC profiles. We investigated whether (1) honey bee workers of different subspecies display different CHC profiles under the same environmental conditions as a consequence of their innate differences; and (2) whether the subspecies-specific differences of the CHC profiles can be interpreted as an adaptation to the environmental conditions of their geographic origin. We hypothesized that foragers should have a more distinct subspecies-specific CHC profile than nurse bees because they are exposed to harsher environmental conditions outside the hive.

## Materials and methods

### Experimental set-up

We performed a common garden experiment, by keeping queen-right colonies of six different honey bee subspecies (*A. m. carnica, A. m. iberiensis, A. m. ligustica, A. m. macedonica, A. m. mellifera*, and *A. m. ruttneri*) in the departmental apiary of the University of Würzburg from April to September 2018. By doing so, we ensure that the colonies experience the same environmental conditions, thus the differences in their CHC composition mainly respond to their genetic differences.

The mated queens came from populations central to the current distribution range of each respective sub-species, namely: Würzburg, Germany: *A. m. carnica*; Bragança, Portugal: *A. m. iberiensis*; Reggio Emilia, Italy: *A. m. ligustica*; Neo Moudania, Greece: *A. m. macedonica*; Malta, Malta: *A. m. ruttneri*. Finally, *A. m. mellifera* queens originated from a protected population in Chimay, Belgium, to avoid analyzing admixed individuals, as this subspecies genetic integrity is threatened in large tracts of its native range by C-derived gene flow [45].

The queens were introduced in queen-less *A. m. carnica* hives for the initial set-up. To avoid drifting of foragers, the hives were placed in pairs of the same subspecies on hive stands, facing the opposite direction, and the hive stands were grouped by subspecies. Furthermore, the hive boxes were sorted in different color combinations, helping the returning foragers to locate their own hive. Thus, intending to reduce the drifting risk within and between the groups of hives [46].

To assure complete turnover in the *A. m. carnica* recipient hives, workers were collected from each colony after a minimum of two months from queen introduction. The workers were immediately killed by immersing them in liquid nitrogen and stored in labeled Eppendorf tubes at -20 °C until hexane extraction. Five workers per task (nursing and foraging) were collected from two different hives of each subspecies, for a total of 20 worker bees per subspecies. Samplings took place between 7:00 a.m. and 12:00 p.m. between July and September 2018. Individual workers inside the hive who poked their heads into an open brood cell for at least 10 seconds were identified as nurse bees. Returning pollen foragers were identified by a considerable pollen load in the corbiculae of their hind legs.

### Extraction of CHCs and chemical analysis

After defrosting, honey bees were immersed in hexane for 10 minutes in 1.5 ml glass vials to extract their CHCs. Subsequently, the solvent was evaporated using a gentle stream of CO_2_, until the total volume was 100 µl to transfer it into a 300 µl glass insert. The insert was placed in a 1.5 ml glass vial. 1 µl of each extract was injected into an Agilent 7890A Series Gas Chromatography System (GC) coupled to an Agilent 5975C Mass Selective Detector (MS) (Agilent Technologies, Waldbronn, Germany). The GC was equipped with a J & W, DB-5 fused silica capillary column (30 m x 0.25 mm ID; df = 0.25 µm; J & W, Folsom, CA, USA). The GC temperature was programmed from 60 to 300 °C with a 5 °C/min heating rate and held for 10 min at 300 °C. Helium was used as carrier gas with a constant flow of 1 ml/min. The injection was carried out at 300 °C in the split-less mode for 1 min with an automatic injector. The electron impact mass spectra (EI-MS) were recorded at 70 eV and with a source temperature of 230 °C.

The chromatograms were analyzed using the data analysis software package ‘MSD ChemStation F.01.00.1903’ for Windows (Agilent Technologies, Waldbronn, Germany). The area of each peak was determined by integration and the initial threshold of the integration parameters was set on 15. Initial area reject was set on 1, initial peak width was set on 0.02 and shoulder detection was off. The compounds were identified by their retention indices and diagnostic ions of their mass spectra. Double bond positions of monounsaturated hydrocarbons were identified by dimethyl disulfide derivatization [47]. Compounds that represented less than 0.01% of the total ion count of a sample and compounds in less than 50% of the extracts in a group were excluded from the analysis to exclude concentration effects and to compare group-specific profiles. The abundances of the compounds were quantified, for each sample, as the proportion (%) that the area of the corresponding peaks represented from the sum of the area of all the peaks included in the analysis.

Additionally, we calculated the relative abundance of the different hydrocarbon classes (i.e. n-alkanes, alkenes, alkadienes, and methyl-branched alkanes), as well as the mean chain length, in the CHC profile of each bee, and compared them between subspecies, for the nurses and the foragers separately. The relative abundances of the different hydrocarbon classes were calculated as the sum of the relative abundances of all the individual compounds of a corresponding class that were found in the CHCs extract of a bee. The mean chain length corresponds to the weighted mean of the chain length of all the compounds found in the CHC profile of each bee, using the relative abundance of the compounds as their weights.

### Climate data

Climatic conditions were characterized by calculating the average temperature (°C) and precipitation (mm) for the countries of origin of the mated queens of each subspecies, using the WorldClim 2.1 [48] downscaled CRU-TS 4.06 data set [49]. Subsequently, we correlated these parameters with the CHC compositions of the subspecies.

### Statistical analysis

The chemical similarity between task and subspecies was visualized via hierarchical cluster analyses, using the mean abundance of each compound for each group, the Bray-Curtis dissimilarity index, and the Wards’ grouping method.

A bi-dimensional non-metric multidimensional scaling (NMDS) was used to visualize the CHC composition differences separately for nurse bees and foragers, using the Bray-Curtis dissimilarity index to estimate the inter-individual chemical dissimilarity.

The difference in the CHC composition between subspecies was tested via a one-way PERMANOVA. The permutations were restricted to the same task-performance groups (i.e. Comparing nurses or foragers between subspecies). Additionally, pairwise comparisons with FDR Benjamini & Hochberg adjustments were performed to test the significance of the differences between subspecies, for the nurses and foragers separately.

The difference in CHC composition variability between nurses and foragers was tested via a one-way permutational test of multivariate homogeneity, with the permutations restricted to the subspecies (i.e. Contrasting the variance in the CHC profiles between nurses and foragers of the same subspecies).

The relative abundance of the different substance classes of hydrocarbons between subspecies was tested, for nurse and forager bees separately, with Kruskal Wallis tests, followed by Dunn’s tests with FDR Benjamini & Hochberg adjustment. The relative abundance of each hydrocarbon substance class was calculated by summing up the relative abundances of all the compounds of different classes in the CHC profile of each honey bee worker.

Likewise, the weighted mean chain length was calculated for every individual CHC profile, and its difference was tested between subspecies, for nurse and forager bees separately, with Kruskal Wallis tests, followed by Dunn’s tests with FDR Benjamini & Hochberg adjustment. The weighted mean chain length was calculated using the relative abundance of the compounds as the weights for averaging their chain length.

The correlation of the average temperature (°C) and precipitation (mm) for the countries of origin of each subspecies with the ratio of olefins (unsaturated hydrocarbons) to n-alkanes, and with the mean chain length of the hydrocarbons, in the CHC profile of the honey bee workers was tested via Spearman’s correlation tests. The olefins to n-alkanes ratio was calculated by summing the relative abundance of the alkenes and alkadienes in the CHC profiles of every bee and dividing it by the respective relative abundance of n-alkanes.

Statistics were performed using R 4.4.0 [50], with the IDE RStudio v2024.4.1.748 [51], and the packages vegan v2.6.6.1 [52], dunn.test v1.3.6 [53], ggdendro v0.2.0 [54], raster v3.6.26 [55], geodata v0.6.2 [56], and tidyverse v2.0.0 [57].

## Results

The average composition of the CHC profiles of the honey bee subspecies differs between nurse bees and foragers, except for *A. m. iberiensis* foragers, which are placed in the same cluster with *A. m. iberiensis and A. m. mellifera* nurses (Figure 1) meaning that the CHC composition of *A. m. iberiensis* foragers is more similar to their and *A. m. mellifera* nurse bees than to foragers of the other subspecies. The characterization of the CHC profiles revealed that, when they are different among *A. mellifera* subspecies, they differ mainly in the relative abundance of their compounds, although, each subspecies displayed one or a few compounds that were not present in the other subspecies (see Supplementary tables S2 and S1). The relative abundance of each substance class differed among subspecies in the CHC profiles of the workers (p-value < 0.05; Kruskal-Wallis test; Table 1), except for the alkenes (p-value > 0.05) in both nurse bees and foragers (Figure 2).

**Table 1:**
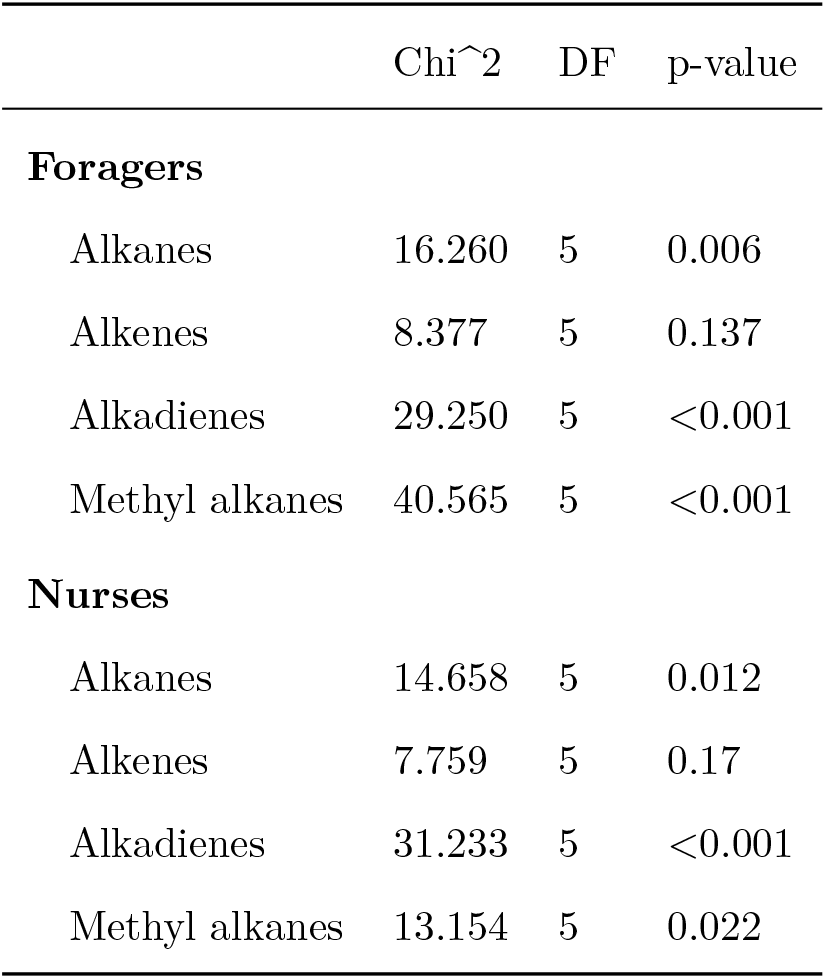
Kruskal-Wallis results for subspecific differences in abundance of hydrocarbon classes in the CHC profile of honey bee workers. Chi2 - χ^2^; DF - degrees of freedom.

**Figure 1.**
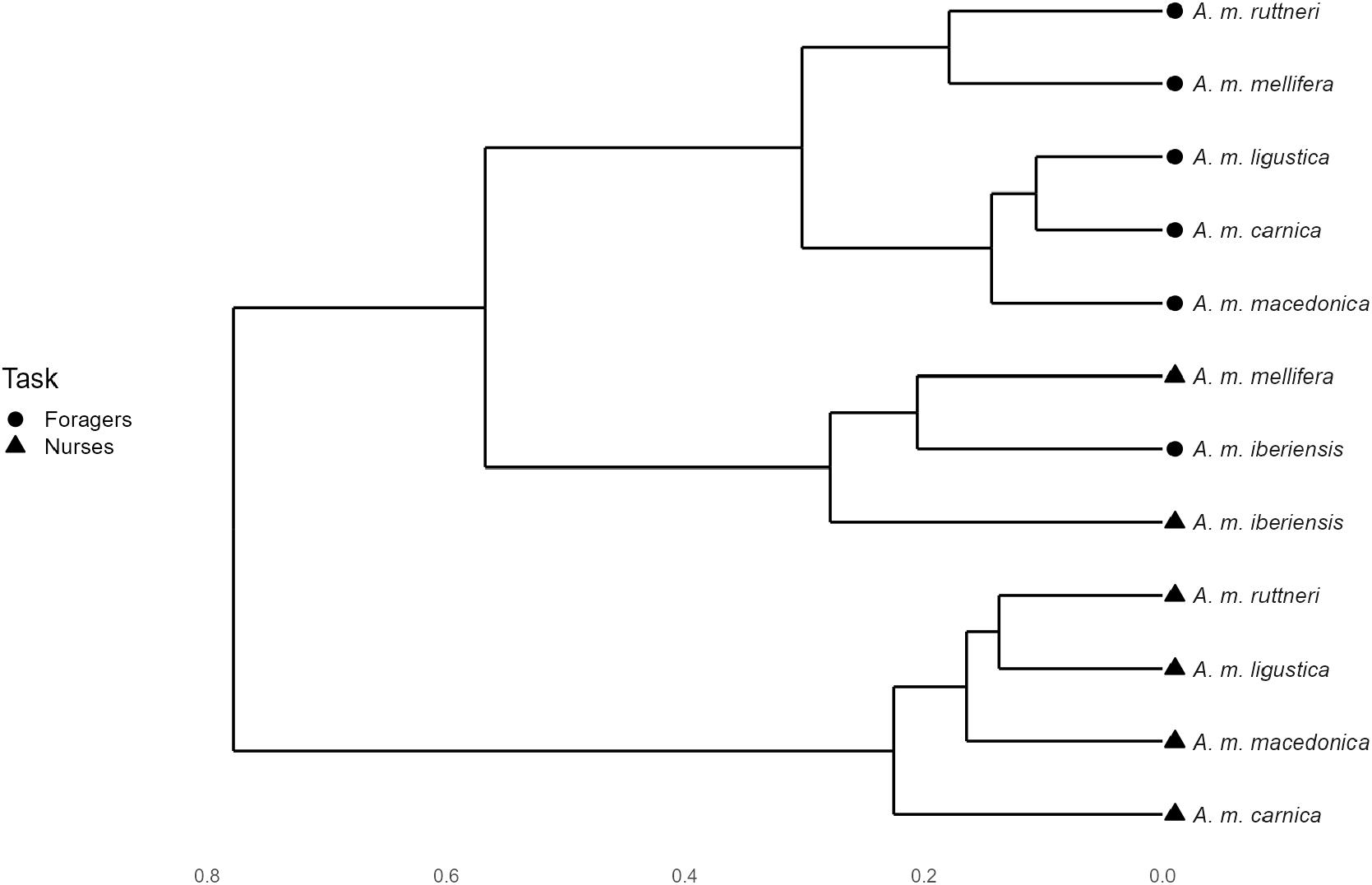
Dendrogram of the CHC profiles hierarchical cluster analysis of *A. mellifera* subspecies worker bees.

**Figure 2.**
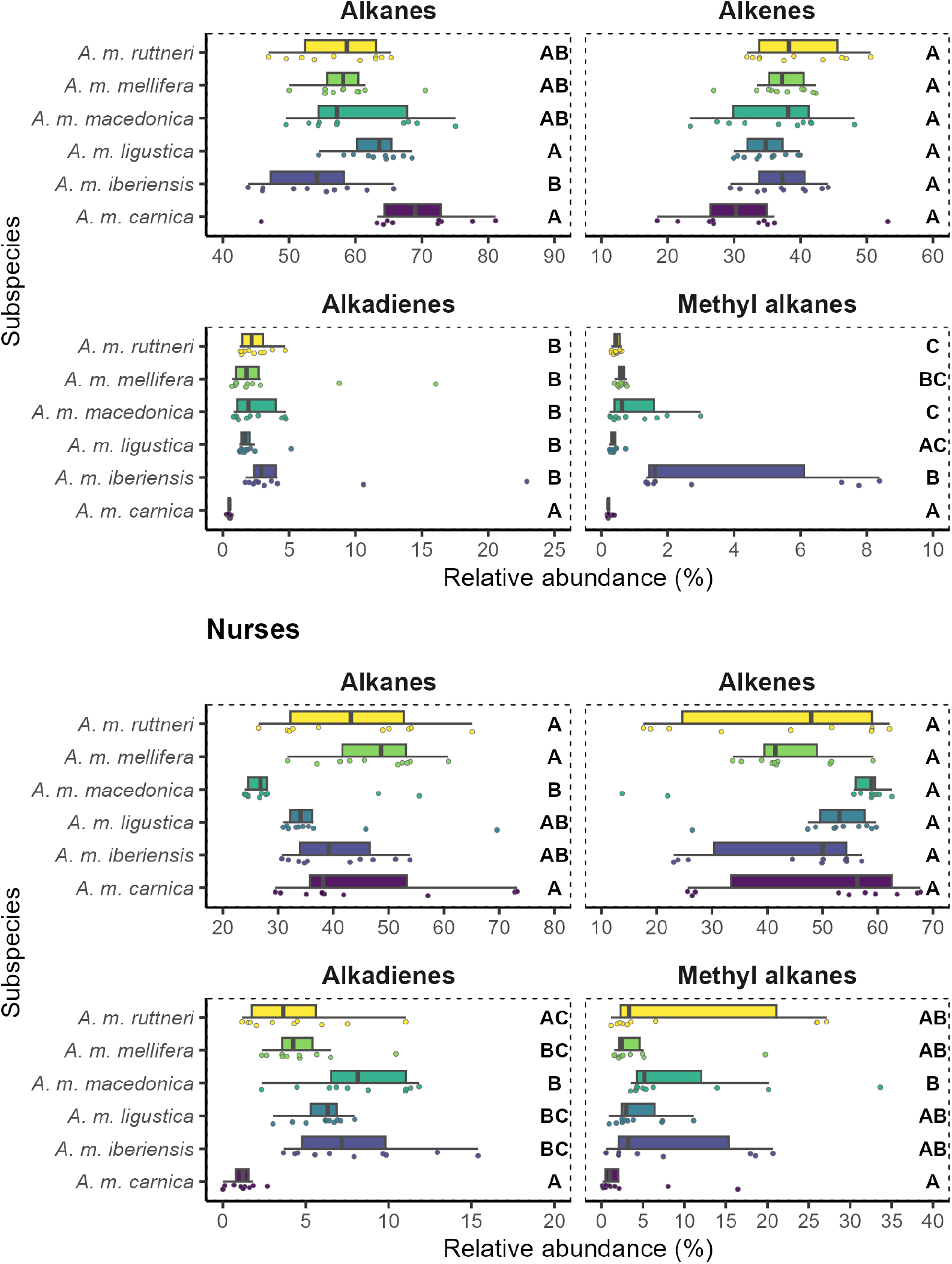
Box plot of the relative abundance of each compound class in the CHC profile of worker bees of different *A. mellifera* subspecies. The figure is divided regarding the task of the worker bees (foragers and nurse bees). Subspecies with significantly different abundance of a hydrocarbon class (p-value < 0.050) are labeled with different letters, within the plot of the corresponding compound class. Significance of pair-wise comparisons was obtained with Dunn’s tests and a FDR Benjamini & Hochberg adjustment, after a Kruskal Wallis tests.

Regarding the foragers, most subspecies differed significantly in their CHC profiles (p-value < 0.05; Figure 3; Tables 2 and 3). Only *A. m. macedonica, A. m. carnica* and *A. m. ligustica* did not differ from each other. Although all other subspecies have a specific CHC profiles, the strongest differences appear between *A. m. carnica* and *A. m. iberienses*, with the former having a higher relative amount of alkanes and the latter having higher relative amounts of methyl-branched hydrocarbons and alkadienes (Figure 2). Interestingly, there was no difference in the weighted mean chain length between the foragers of the studied subspecies (p-value > 0.05; Figure 4; Table 4).

**Table 2:** One-way PERMANOVA contrasting the CHC composition between subspecies. The permutations (n=1000) were restricted to the task-performance group. Df - degrees of freedom; SS - sum of squares; R2 - R squared; F - F-value; SES - standard effect size.

|  | Df | SS | R2 | F | SES | p-value |
| --- | --- | --- | --- | --- | --- | --- |
| Subspecies | 5 | 2.185 | 0.203 | 5.795 | 25.803 | 0.001 |
| Residual | 114 | 8.596 | 0.797 |  |  |  |
| Total | 119 | 10.781 | 1.000 |  |  |  |

**Table 3:**
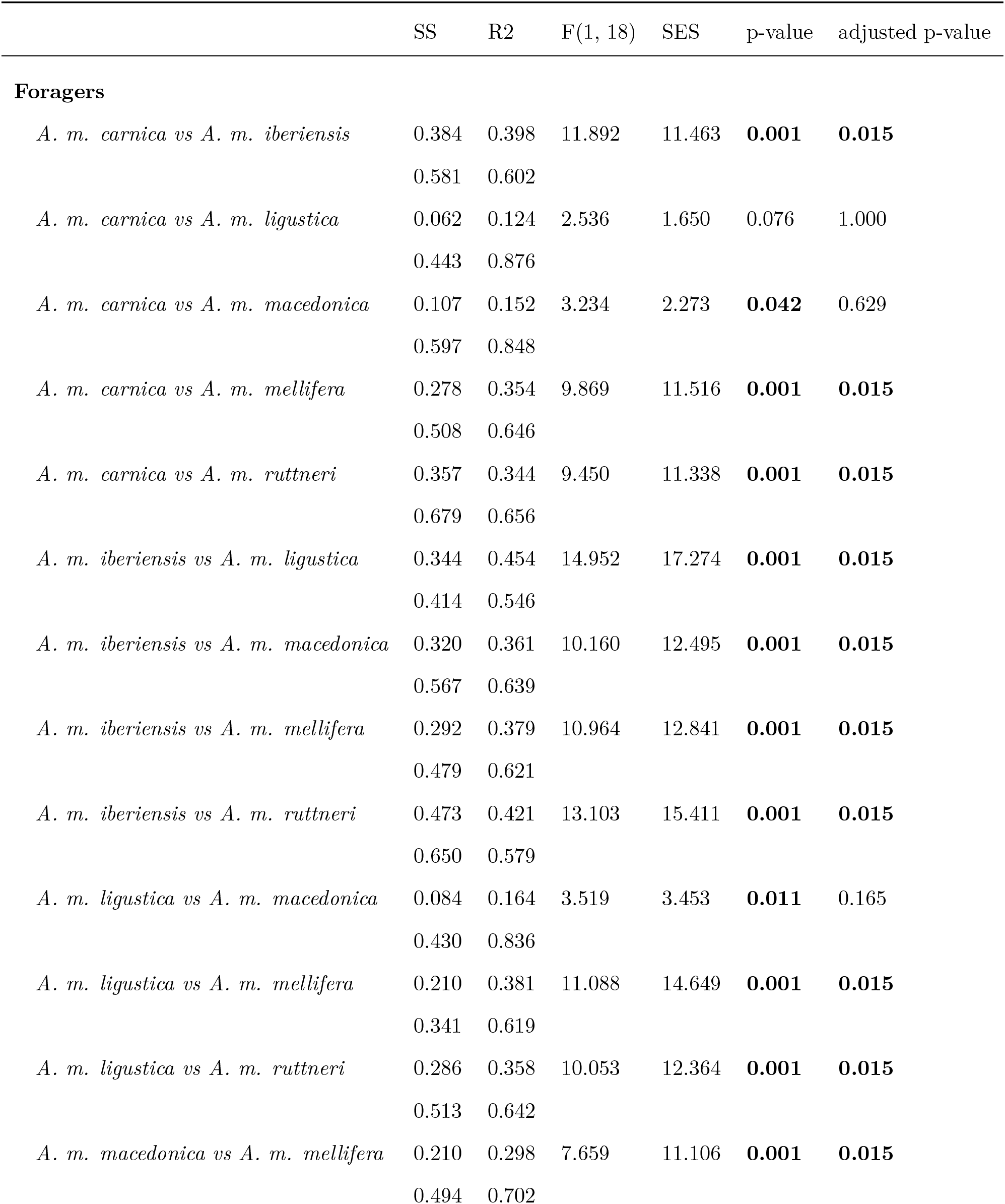
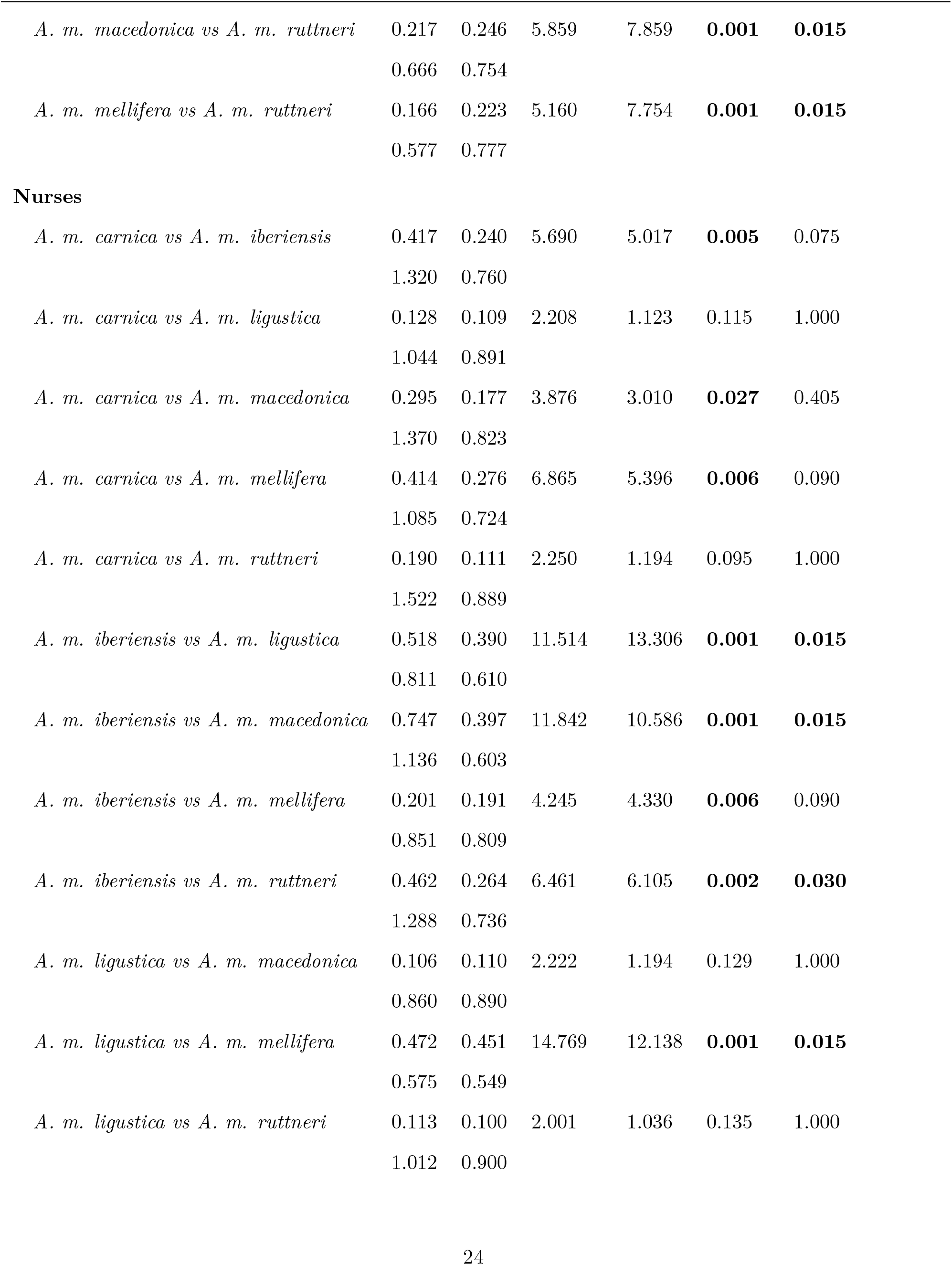
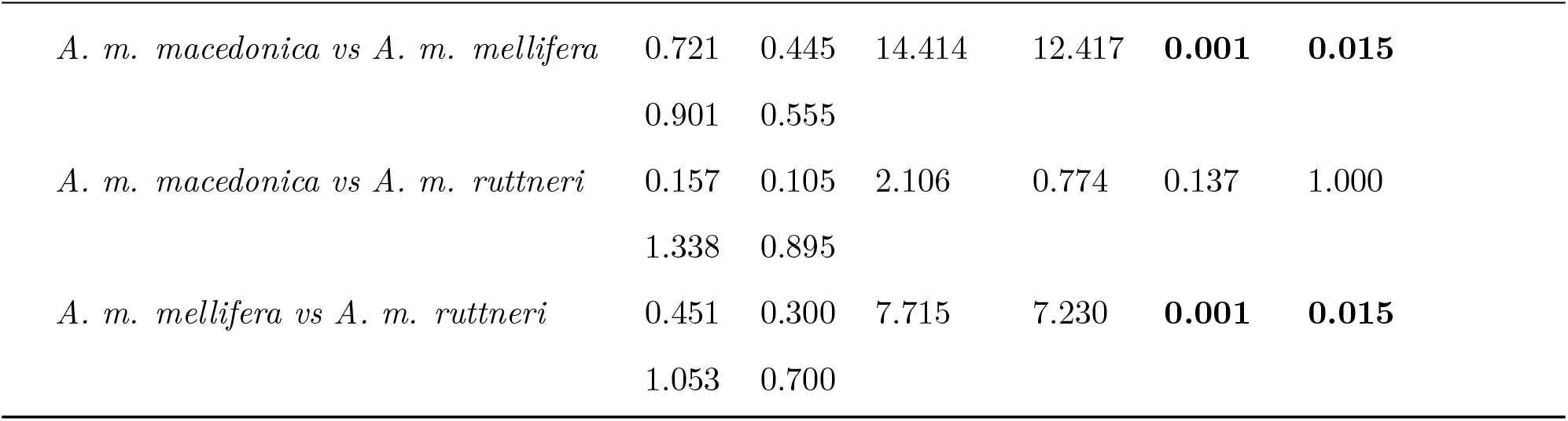
Pairwise PERMANOVA results contrasting the CHC composition between honey bee subspecies for the nurses and foragers separately. SS - sum of squares; R2 - R squared; F - F-value, SES - standard effect size. In the case of the SS and R2 the value for each comparison is displayed over that of the corresponding residuals. The adjusted p-values result from a FDR Benjamini & Hochberg adjustment. p-values < 0.05 are marked in bold.

**Table 4:** Kruskal-Wallis results for subspecific difference in mean chain length of hydrocarbons in the CHC profile of honey bee workers. Chi^2^ - χ^2^; DF-degrees of freedom.

|  | Chi2 | Df | p-value |
| --- | --- | --- | --- |
| Foragers | 12.955 | 5 | 0.024 |
| Nurses | 19.965 | 5 | 0.001 |

**Figure 3.**
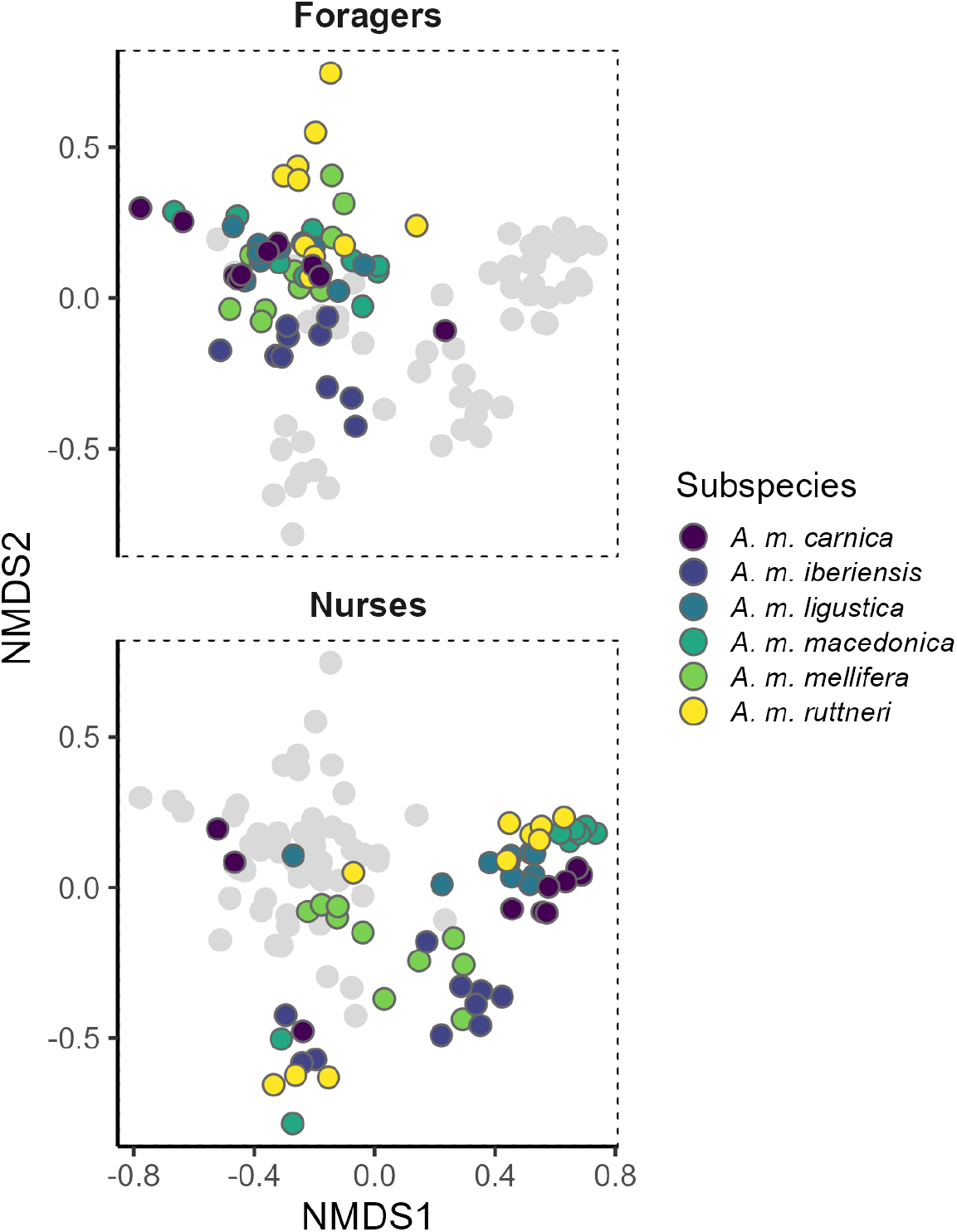
Bi-dimensional Non-metric Multidimensional Scaling analysis on a dissimilarity matrix of the CHC profile of honey bee workers. Stress value: 0.140. The figure is divided in two sections regarding the task of the worker bees (Foragers on the top, and nurse bees at the bottom). Both sections show the same NMDS plot, but only the points representing bees of the corresponding task are colored, while the others are shown in grey.

**Figure 4.**
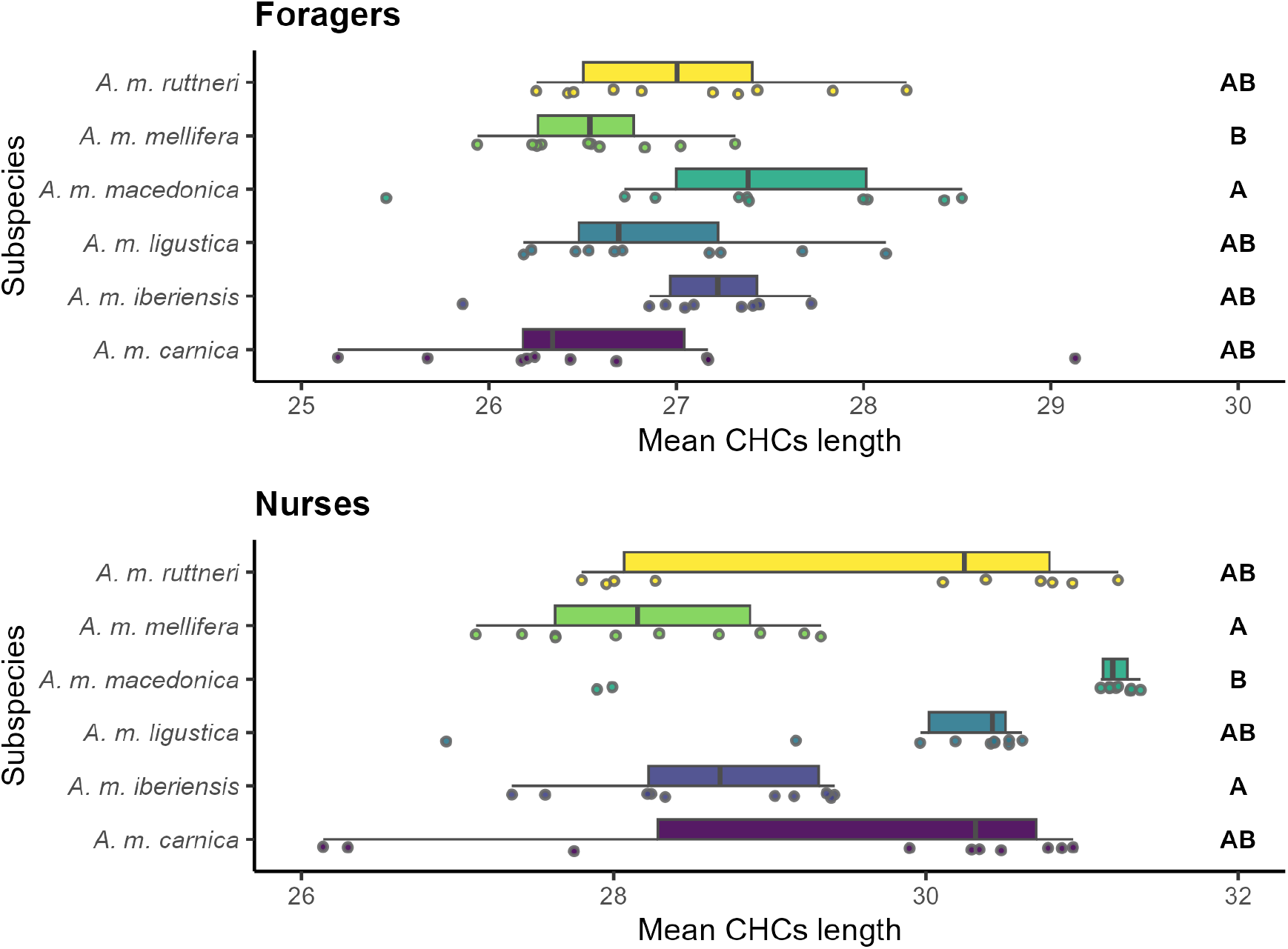
Box plot of the weighted mean chain length of hydrocarbons in the CHC profile of worker bees of different *A. mellifera* subspecies. The figure is divided regarding the task of the worker bees (foragers and nurse bees). Subspecies that significantly differ (p-value < 0.050) are labeled with different letters. Significance of pair-wise comparisons was obtained with Dunn’s tests and a FDR Benjamini & Hochberg adjustment, after a Kruskal Wallis tests.

As expected, there were fewer differences in the CHC profiles among subspecies in nurse bees than in foragers (p-value < 0.05; Figure 3; Tables 2 and 3). Regarding the nurse bees *A. m. mellifera* and *A. m. iberiensis* were not different from each other, but they were different from all the other subspecies. There were no differences among *A. m. macedonica, A. m. carnica, A. m. ligustica*, and *A. m. mellifera*. Of all subspecies, *A. m. macedonica* has the lowest relative amounts of alkanes in favor of higher amounts of methyl-branched alkanes and alkadienes. In contrast, *A. m. carnica* shows relatively low amounts of methyl-branched alkanes and alkadienes (Figure 2).

There was more variation in the weighted mean chain length of CHC profiles in nurse bees than foragers with the highest weighted mean chain length in *A. m. macedonica* and the lowest weighted mean chain length in *A. m. mellifera* and *A. m. iberiensis* nurse bees (Figure 4). In addition, the CHC profile of nurse bees occupied a significantly larger chemical space than that of the foragers (p-value < 0.05; Figure 3; Supplementary Table S3).

The average temperature for the country of origin of the subspecies correlated positively with the ratio of unsaturated hydrocarbons to n-alkanes only in the nurse bees, while no correlation was observed for the foragers. In contrast, the average temperature correlated negatively with the weighted mean chain length of the CHC profiles in the foragers, but they did not correlate for the case of the nurse bees (Supplementary Figure S1).

The average precipitation for the country of origin of the subspecies correlated negatively with the ratio of unsaturated hydrocarbons to n-alkanes in the nurse bees and with the weighted mean chain length in the foragers (Supplementary Figure S2). No correlation was observed for the mean precipitation and the unsaturated to n-alkanes ratio in the foragers, nor with the weighted mean chain length of the nurse bees. The correlations with the climate parameters and the CHC profiles followed the prediction of adapting to drought stress only in nurse bees but not in foragers. Overall, the correlation coefficients were low (R < 0.4, Spearman’s test).

## Discussion

In the present study, we investigated (1) whether honey bee subspecies exhibit a differentiated CHC profile, and (2) whether potential CHC differences between subspecies can be explained by adaptation to climatic conditions or by genetic drift. We show that the composition of CHC profiles of honey bee workers differ moderately between subspecies with more pronounced differences between foragers than nurses. Keeping the subspecies under the same environmental conditions suggests that the subspecies’ differences in CHC profiles have a genetic basis. The cluster analysis reveals similarities among honey bee subspecies belonging to the same evolutionary lineage. In nurse bees, the members of the lineage M, *A. m. mellifera* and *A. m. iberiensis*, and the members of the lineage C, *A. m. carnica, A. m. ligustica*, and *A. m. macedonica*, are more similar among each other than between the different lineages [6, 7, 58, 59]. This indicates phylogenetic constraint as a factor for CHC profile diversification. However, an effect of genetic drift on the CHC profiles of the subspecies cannot be ruled out. The only exception is the A-lineage *A. m. ruttneri*, which clusters with the members of the lineage C, but this subspecies is highly admixed with C-derived *A. m. ligustica* bees [6, 58, 59]. This contrasts to CHC profiles in ants, which have been shown to evolve independently from phylogeny, allowing them to rapidly adapt their CHC profiles to environmental selection pressures [60]. In foragers, the members of the lineages do not cluster together with the exception of lineage C, suggesting an impact of environmental factors on their CHC profiles.

In contrast to our expectations, we did not find a strong adaptation of the subspecies CHC profiles to their climatic conditions. We had expected that particularly foragers from environments with more drought stress (e.g. *A. m. iberiensis*) would harden their CHC profile by increasing the ratio of unsaturated hydrocarbons to n-alkanes and the weighted mean chain length. However, the correlation of these CHC profile parameters with temperature and precipitation supported the adaptation hypothesis only in the nurse bees but not in foragers and, in addition, was very weak. Thus, we suggest that the evolution of *A. mellifera* subspecies-specific CHC profiles has been under a stronger influence of phylogenetic constraint and/or genetic drift than climatic conditions.

By performing a common garden experiment with all investigated subspecies, we ensured that all colonies were kept under the same environmental conditions. Thus, the differences in the CHC composition among subspecies should be due to genetic differences. Although, there were no consistent signs of adaptation in the CHC profiles of the subspecies to the climatic conditions in their country of origin, we found less variation in the CHC profiles between individual foragers within a subspecies compared to nurse bees of the same subspecies among all investigated honey bee subspecies. This might be due to an overall stronger desiccation stress on foragers, independent of their local environment. Foragers might be restricted from occupying a large chemical space, as we see in nurse bees, because their CHC profile is already adapted to a harsher environment outside the hive. Alternatively, the smaller chemical space found in foragers is an adaptation to nestmate recognition helping the guard bees to better discriminate between nestmates and non-nestmates. To test this hypothesis, a future study should compare foragers and nurse bees from different honey bee colonies.

Honey bee workers perform several tasks inside and outside the nest during their lifetime. The task performance of each worker is influenced by its age, sucrose responsiveness, genotype, environmental conditions, and previous experience [61–67]. The most pronounced differences in behaviour and physiology occur between nurse bees and foragers. While nurse bees provide brood and queen with food, the foragers leave the hive to collect pollen, nectar, water or propolis [68]. We find that these differences between nurse bees and foragers correlate with differential CHC profiles across subspecies. This could be a result of their different environments. While nurse bees normally do not leave the hive, foragers leave the colony daily for up to several hours. Our findings directly support data from other social insects performing tasks outside and inside the colony [38, 39]. Generally, it has been hypothesized that the differences in the CHC composition of foragers compared to nurse bees could be a response to the harsher environment that foragers face outside the hive [31]. This is supported by the fact that foragers display a higher abundance of alkanes and a lower abundance of alkenes than nurse bees. This shift in alkane-to-alkene ratio makes the CHC profile of the foragers more waterproof than that of nurse bees [23, 24, 41, 42, 69]. In contrast, foragers display hydrocarbons with shorter mean chain lengths than nurses, which weakens the waterproofing of the foragers CHC profile and thus contradicts a simple response of the CHC profiles to stronger desiccation stress in foragers. Although the shift in abundance of the different substance classes is likely to have a stronger effect on the waterproofing of the CHC layer than a shift in mean chain length, other factors, such as intraspecific chemical signaling using hydrocarbons, might shape the CHC profiles [23, 26, 29, 41, 41, 42, 70, 71]. Part of the variation in the CHC profile across nurses and foragers might allow an individual worker to assess the task of an encountered individual in the nest.

## Conclusions

Overall, some *Apis mellifera* subspecies show a specific CHC profile when comparing the nurse and forager profiles separately from each other. In addition, our study reveals a very strong shift in the CHC profiles from nurses to foragers in all investigated subspecies. We hypothesize that these CHC differences between subspecies evolved in parallel to morphological traits within these populations due to allopatry. Although there were no signs of adaptation of subspecific CHC profiles to their native climatic conditions, the adaptation of forager CHC profiles of all subspecies compared to the nurse bee profiles was evident. Thus, we hypothesize that the more restricted chemical space of forager CHC profiles compared to the chemical space of nurse bees is an adaptation to harsher climatic conditions outside of the hive in all subspecies.

## Acknowledgements

We would like to express our gratitude to Dirk Ahrens-Lagast, who provided the *A. m. carnica* queens used for this study.

## Data availability

The data and code used for this study are not included in this manuscript, but will be made publicly available as supplementary material for the corresponding paper at the time of publication.

## Supplementary material

**Table S1:**
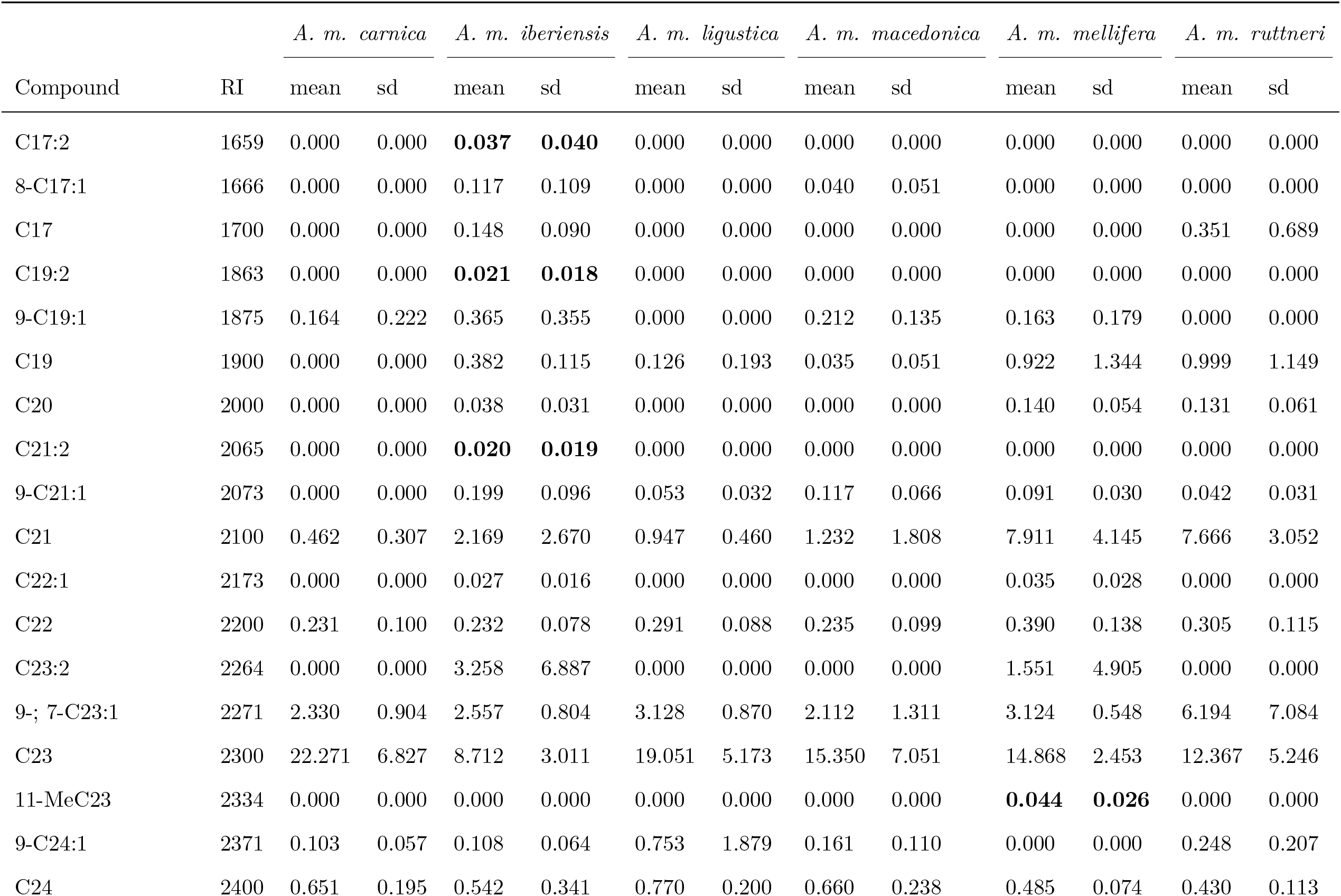
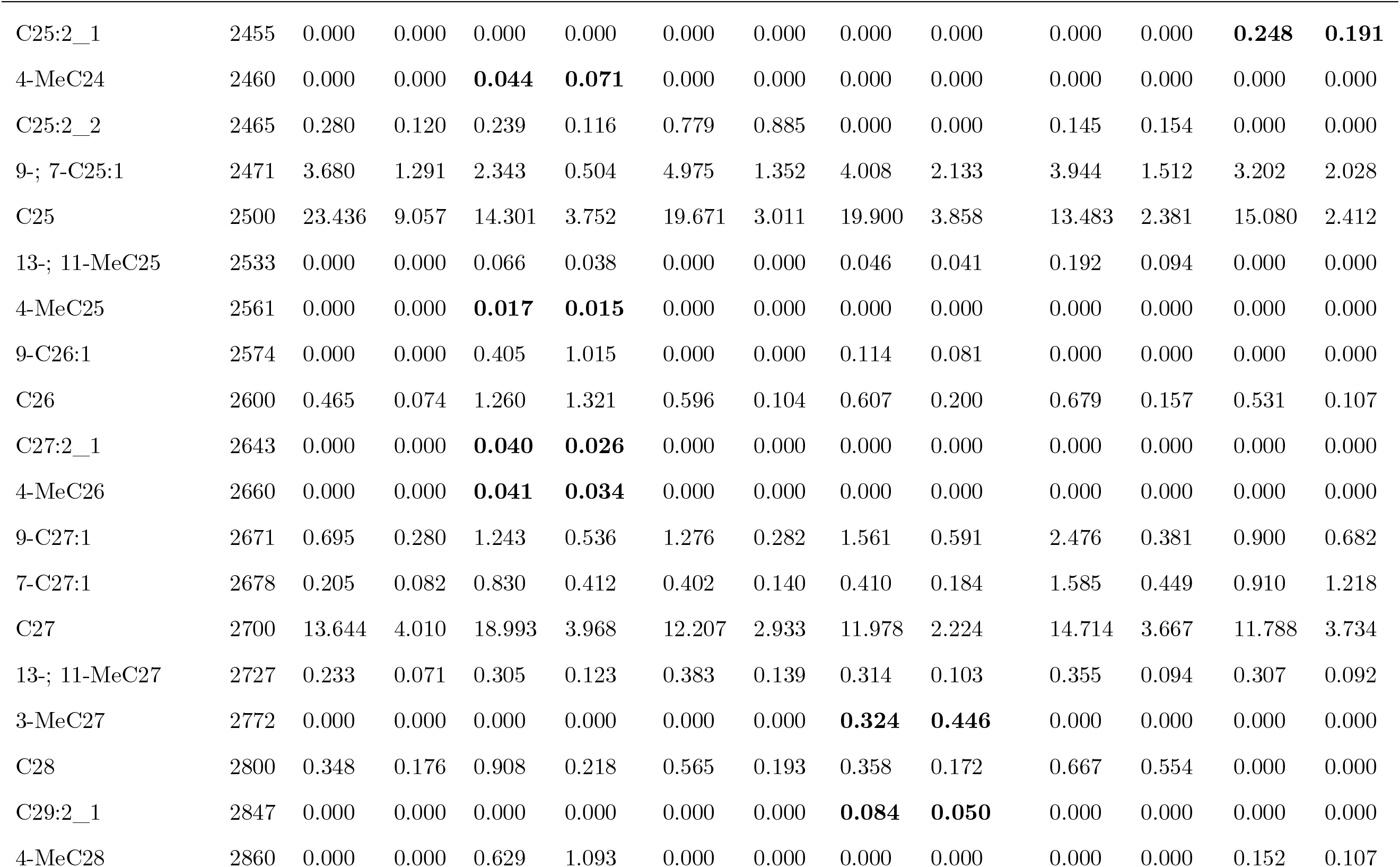
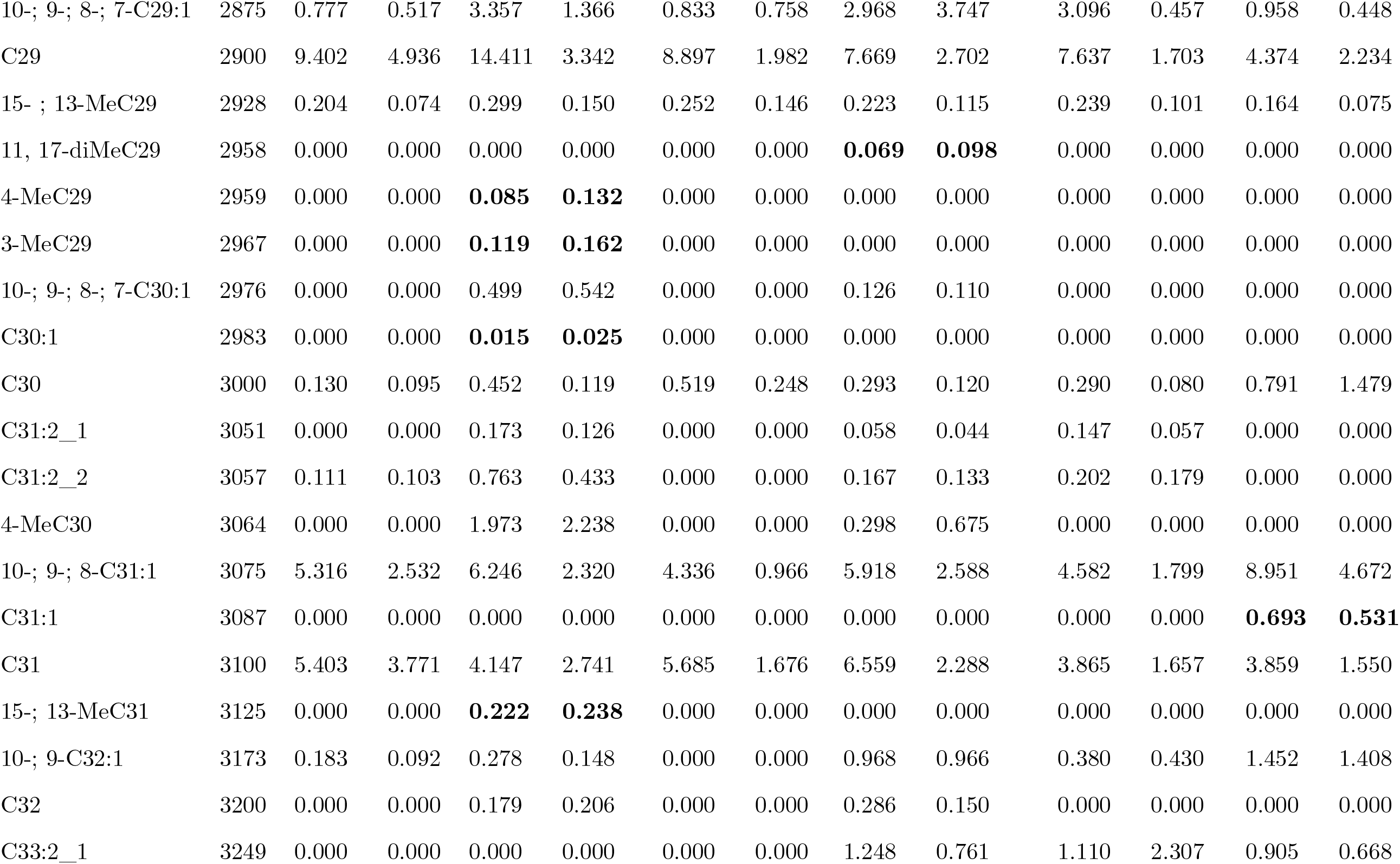
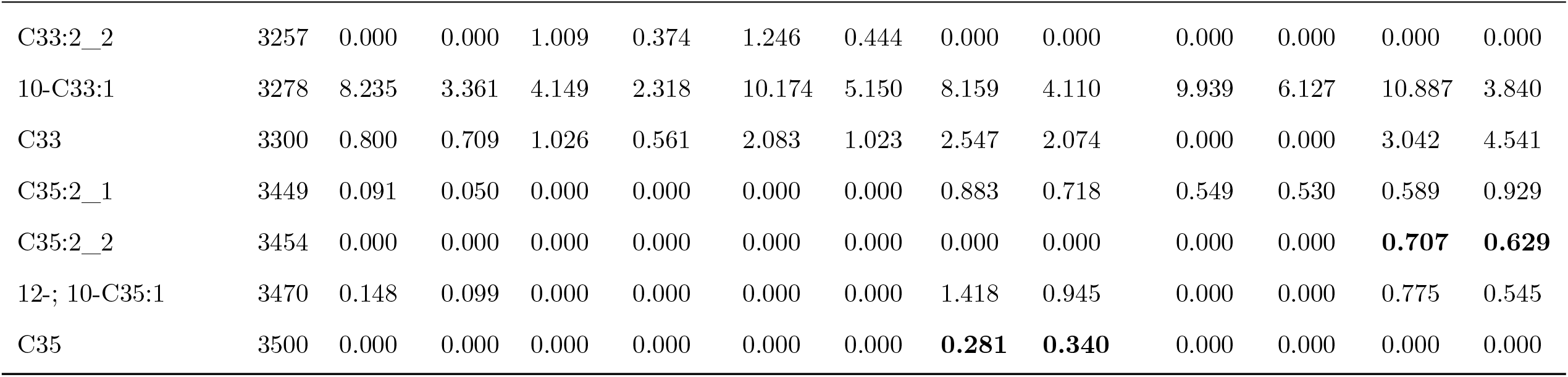
CHC profile of forager bees from different *A. mellifera* subspecies showing mean relative abundance (%) and standard deviation (sd) of all compounds. Mean and sd values are in bold for the cases a compound is present in only one subspecies.

**Table S2:** CHC profile of nurse bees from different *A. mellifera* subspecies showing mean relative abundance (%) and standard deviation (sd) of all compounds. Mean and sd values are in bold for the cases a compound is present in only one subspecies.

| Compound | RI | *A. m. carnica* |  | *A. m. iberiensis* |  | *A. m. ligustica* |  | *A. m. macedonica* |  | *A. m. mellifera* |  | *A. m. ruttneri* |  |
| --- | --- | --- | --- | --- | --- | --- | --- | --- | --- | --- | --- | --- | --- |
|  |  | mean | sd | mean | sd | mean | sd | mean | sd | mean | sd | mean | sd |
| C17:2 | 1659 | 0.000 | 0.000 | **0.048** | **0.057** | 0.000 | 0.000 | 0.000 | 0.000 | 0.000 | 0.000 | 0.000 | 0.000 |
| 8-C17:1 | 1666 | 0.000 | 0.000 | 0.178 | 0.182 | 0.000 | 0.000 | 0.000 | 0.000 | 0.020 | 0.028 | 0.000 | 0.000 |
| C17 | 1700 | 0.000 | 0.000 | 0.277 | 0.388 | 0.000 | 0.000 | 0.000 | 0.000 | 0.028 | 0.026 | 0.000 | 0.000 |
| C19:2 | 1863 | 0.000 | 0.000 | **0.019** | **0.023** | 0.000 | 0.000 | 0.000 | 0.000 | 0.000 | 0.000 | 0.000 | 0.000 |
| 9-C19:1 | 1875 | 0.000 | 0.000 | 0.243 | 0.124 | 0.230 | 0.159 | 0.180 | 0.199 | 0.281 | 0.213 | 0.053 | 0.042 |
| C19 | 1900 | 0.000 | 0.000 | 0.943 | 0.989 | 0.137 | 0.101 | 0.068 | 0.090 | 0.228 | 0.197 | 0.131 | 0.073 |

Table S2: (continued)
| Compound | RI | *A. m. carnica* |  | *A. m. iberiensis* |  | *A. m. ligustica* |  | *A. m. macedonica* |  | *A. m. mellifera* |  | *A. m. ruttneri* |  |
| --- | --- | --- | --- | --- | --- | --- | --- | --- | --- | --- | --- | --- | --- |
|  |  | mean | sd | mean | sd | mean | sd | mean | sd | mean | sd | mean | sd |
| C20 | 2000 | 0.000 | 0.000 | 0.037 | 0.039 | 0.000 | 0.000 | 0.000 | 0.000 | 0.027 | 0.013 | 0.000 | 0.000 |
| 9-C21:1 | 2073 | 0.000 | 0.000 | 0.115 | 0.101 | 0.000 | 0.000 | 0.000 | 0.000 | 0.046 | 0.027 | 0.000 | 0.000 |
| C21 | 2100 | 0.000 | 0.000 | 2.137 | 1.301 | 1.426 | 1.186 | 0.626 | 0.735 | 1.118 | 0.528 | 1.206 | 0.689 |
| C22:1 | 2173 | 0.000 | 0.000 | 0.000 | 0.000 | 0.000 | 0.000 | 0.000 | 0.000 | **0.013** | **0.016** | 0.000 | 0.000 |
| C22 | 2200 | 0.000 | 0.000 | 0.221 | 0.098 | 0.151 | 0.052 | 0.086 | 0.079 | 0.270 | 0.086 | 0.217 | 0.114 |
| C23:2 | 2264 | 0.000 | 0.000 | 0.123 | 0.173 | 0.000 | 0.000 | 0.000 | 0.000 | 0.129 | 0.176 | 0.000 | 0.000 |
| 9-; 7-C23:1 | 2271 | 0.555 | 0.736 | 1.010 | 0.538 | 0.802 | 0.736 | 0.599 | 0.501 | 1.890 | 0.880 | 1.181 | 0.276 |
| C23 | 2300 | 7.740 | 9.622 | 5.336 | 4.512 | 4.522 | 4.518 | 3.625 | 4.816 | 7.146 | 4.388 | 5.750 | 4.269 |
| 11-MeC23 | 2334 | 0.000 | 0.000 | 0.061 | 0.078 | 0.000 | 0.000 | 0.032 | 0.057 | 0.042 | 0.044 | 0.000 | 0.000 |
| 9-C24:1 | 2371 | 0.000 | 0.000 | 0.000 | 0.000 | 0.000 | 0.000 | 0.000 | 0.000 | **0.107** | **0.066** | 0.000 | 0.000 |
| C24 | 2400 | 0.000 | 0.000 | 0.269 | 0.068 | 0.236 | 0.128 | 0.069 | 0.028 | 0.452 | 0.120 | 0.284 | 0.141 |
| C25:2_1 | 2455 | 0.000 | 0.000 | 0.171 | 0.265 | 0.012 | 0.015 | 0.073 | 0.133 | 0.035 | 0.071 | 0.083 | 0.124 |
| 4-MeC24 | 2460 | 0.000 | 0.000 | 0.000 | 0.000 | 0.000 | 0.000 | 0.000 | 0.000 | **0.018** | **0.009** | 0.000 | 0.000 |
| C25:2_2 | 2465 | 0.000 | 0.000 | 0.043 | 0.038 | 0.078 | 0.157 | 0.000 | 0.000 | 0.105 | 0.076 | 0.000 | 0.000 |
| 9-; 7-C25:1 | 2471 | 1.269 | 1.941 | 0.722 | 0.345 | 1.091 | 1.251 | 0.549 | 0.602 | 1.735 | 1.209 | 0.599 | 0.632 |
| C25 | 2500 | 7.888 | 10.901 | 4.160 | 1.682 | 6.135 | 4.483 | 2.783 | 0.836 | 9.359 | 4.760 | 5.967 | 4.708 |
| 13-; 11-MeC25 | 2533 | 0.175 | 0.315 | 0.355 | 0.415 | 0.142 | 0.057 | 0.304 | 0.529 | 0.255 | 0.327 | 0.358 | 0.359 |
| 5-MeC25 | 2547 | 0.000 | 0.000 | 0.031 | 0.043 | 0.000 | 0.000 | 0.000 | 0.000 | 0.000 | 0.000 | 0.033 | 0.033 |
| C26:1_1 | 2571 | 0.000 | 0.000 | 0.000 | 0.000 | 0.000 | 0.000 | 0.000 | 0.000 | **0.078** | **0.033** | 0.000 | 0.000 |

Table S2: (continued)
| Compound | RI | *A. m. carnica* |  | *A. m. iberiensis* |  | *A. m. ligustica* |  | *A. m. macedonica* |  | *A. m. mellifera* |  | *A. m. ruttneri* |  |
| --- | --- | --- | --- | --- | --- | --- | --- | --- | --- | --- | --- | --- | --- |
|  |  | mean | sd | mean | sd | mean | sd | mean | sd | mean | sd | mean | sd |
| 3-MeC25 | 2571 | 0.000 | 0.000 | 0.028 | 0.024 | 0.010 | 0.013 | 0.027 | 0.032 | 0.000 | 0.000 | 0.042 | 0.029 |
| 9-C26:1 | 2574 | 0.000 | 0.000 | 0.000 | 0.000 | 0.000 | 0.000 | 0.000 | 0.000 | **0.092** | **0.056** | 0.000 | 0.000 |
| 5, 15-diMeC25 | 2583 | 0.000 | 0.000 | 0.000 | 0.000 | 0.000 | 0.000 | **0.061** | **0.102** | 0.000 | 0.000 | 0.000 | 0.000 |
| C26 | 2600 | 0.190 | 0.191 | 0.581 | 0.117 | 0.366 | 0.080 | 0.236 | 0.059 | 0.837 | 0.108 | 0.478 | 0.184 |
| 13-MeC26 | 2630 | 0.000 | 0.000 | 0.078 | 0.080 | 0.039 | 0.011 | 0.077 | 0.094 | 0.060 | 0.063 | 0.077 | 0.086 |
| C27:2_1 | 2643 | 0.000 | 0.000 | 0.230 | 0.378 | 0.023 | 0.024 | 0.059 | 0.105 | 0.039 | 0.050 | 0.000 | 0.000 |
| C27:2_2 | 2649 | 0.000 | 0.000 | 0.000 | 0.000 | 0.000 | 0.000 | 0.000 | 0.000 | **0.023** | **0.027** | 0.000 | 0.000 |
| 4-MeC26 | 2660 | 0.000 | 0.000 | **0.012** | **0.019** | 0.000 | 0.000 | 0.000 | 0.000 | 0.000 | 0.000 | 0.000 | 0.000 |
| 9-C27:1 | 2671 | 0.201 | 0.268 | 0.343 | 0.071 | 0.350 | 0.258 | 0.238 | 0.114 | 1.519 | 0.933 | 0.351 | 0.314 |
| 7-C27:1 | 2678 | 0.055 | 0.077 | 0.314 | 0.157 | 0.108 | 0.077 | 0.054 | 0.016 | 0.933 | 0.787 | 0.098 | 0.068 |
| C27 | 2700 | 12.933 | 4.429 | 15.776 | 3.668 | 10.761 | 1.993 | 10.104 | 3.531 | 19.477 | 3.589 | 13.316 | 3.680 |
| 13-, 11-MeC27 | 2727 | 1.866 | 3.246 | 2.889 | 3.679 | 1.209 | 0.910 | 2.027 | 3.295 | 1.420 | 2.345 | 3.477 | 4.438 |
| 7-MeC27 | 2738 | 0.000 | 0.000 | 0.118 | 0.150 | 0.000 | 0.000 | 0.087 | 0.132 | 0.000 | 0.000 | 0.000 | 0.000 |
| 5-MeC27 | 2747 | 0.000 | 0.000 | 0.047 | 0.063 | 0.000 | 0.000 | 0.045 | 0.065 | 0.000 | 0.000 | 0.062 | 0.080 |
| 11, 15-DiMeC27 | 2756 | 0.000 | 0.000 | 0.121 | 0.226 | 0.000 | 0.000 | 0.112 | 0.160 | 0.000 | 0.000 | 0.000 | 0.000 |
| x, x-DiMeC27 | 2758 | 0.000 | 0.000 | 0.000 | 0.000 | 0.000 | 0.000 | **0.086** | **0.124** | 0.000 | 0.000 | 0.000 | 0.000 |
| y, y-diMeC27 | 2762 | 0.000 | 0.000 | 0.000 | 0.000 | 0.000 | 0.000 | **0.084** | **0.122** | 0.000 | 0.000 | 0.000 | 0.000 |
| z, z-diMeC27 | 2770 | 0.000 | 0.000 | 0.000 | 0.000 | 0.000 | 0.000 | **0.118** | **0.155** | 0.000 | 0.000 | 0.000 | 0.000 |
| C28:1_1 | 2772 | 0.000 | 0.000 | 0.151 | 0.131 | 0.000 | 0.000 | 0.000 | 0.000 | 0.115 | 0.065 | 0.000 | 0.000 |

Table S2: (continued)
| Compound | RI | *A. m. carnica* |  | *A. m. iberiensis* |  | *A. m. ligustica* |  | *A. m. macedonica* |  | *A. m. mellifera* |  | *A. m. ruttneri* |  |
| --- | --- | --- | --- | --- | --- | --- | --- | --- | --- | --- | --- | --- | --- |
|  |  | mean | sd | mean | sd | mean | sd | mean | sd | mean | sd | mean | sd |
| 3-MeC27 | 2772 | 0.000 | 0.000 | 0.000 | 0.000 | 0.026 | 0.034 | 0.000 | 0.000 | 0.000 | 0.000 | 0.021 | 0.026 |
| C28:1_3 | 2781 | 0.000 | 0.000 | 0.288 | 0.324 | 0.000 | 0.000 | 0.000 | 0.000 | 0.156 | 0.065 | 0.000 | 0.000 |
| 5, 17-diMeC27 | 2780 | 0.000 | 0.000 | 0.000 | 0.000 | 0.000 | 0.000 | **0.118** | **0.115** | 0.000 | 0.000 | 0.000 | 0.000 |
| C28 | 2800 | 0.421 | 0.207 | 1.136 | 0.236 | 0.525 | 0.123 | 0.459 | 0.216 | 0.699 | 0.130 | 0.579 | 0.353 |
| 14-; 13-MeC28 | 2830 | 0.000 | 0.000 | 0.335 | 0.451 | 0.116 | 0.115 | 0.300 | 0.415 | 0.129 | 0.255 | 0.438 | 0.641 |
| C29:2_1 | 2847 | 0.000 | 0.000 | 0.025 | 0.020 | 0.000 | 0.000 | 0.000 | 0.000 | 0.038 | 0.031 | 0.000 | 0.000 |
| C29:2_2 | 2853 | 0.000 | 0.000 | 0.000 | 0.000 | 0.000 | 0.000 | 0.000 | 0.000 | **0.136** | **0.086** | 0.000 | 0.000 |
| C29:2_3 | 2860 | 0.000 | 0.000 | 0.120 | 0.054 | 0.000 | 0.000 | 0.000 | 0.000 | 0.114 | 0.027 | 0.000 | 0.000 |
| 4-MeC28 | 2860 | 0.000 | 0.000 | 0.000 | 0.000 | 0.000 | 0.000 | 0.045 | 0.039 | 0.000 | 0.000 | 0.087 | 0.068 |
| 10-; 9-; 8-; 7-C29:1 | 2875 | 0.631 | 0.185 | 4.869 | 2.605 | 0.855 | 0.398 | 0.740 | 0.131 | 4.192 | 1.095 | 0.680 | 0.108 |
| C29 | 2900 | 13.658 | 4.739 | 16.966 | 5.840 | 9.602 | 1.511 | 7.074 | 0.990 | 11.856 | 3.424 | 6.837 | 2.622 |
| 15- ; 13-MeC29 | 2928 | 2.069 | 3.428 | 1.294 | 3.195 | 1.622 | 1.500 | 3.093 | 4.182 | 0.514 | 0.311 | 4.580 | 6.964 |
| 13, 17-diMeC29 | 2951 | 0.000 | 0.000 | 0.000 | 0.000 | 0.000 | 0.000 | 0.195 | 0.252 | 0.086 | 0.102 | 0.000 | 0.000 |
| 11, 17-diMeC29 | 2958 | 0.000 | 0.000 | 0.000 | 0.000 | 0.000 | 0.000 | 0.414 | 0.516 | 0.202 | 0.276 | 0.000 | 0.000 |
| C30:2 | 2959 | 0.000 | 0.000 | **0.626** | **0.620** | 0.000 | 0.000 | 0.000 | 0.000 | 0.000 | 0.000 | 0.000 | 0.000 |
| 10-; 9-; 8-; 7-C30:1 | 2976 | 0.000 | 0.000 | 0.146 | 0.130 | 0.118 | 0.067 | 0.246 | 0.083 | 0.391 | 0.227 | 0.000 | 0.000 |
| 5-,17-; 5,15-diMeC29 | 2980 | 0.000 | 0.000 | 0.000 | 0.000 | 0.000 | 0.000 | 0.000 | 0.000 | **0.058** | **0.106** | 0.000 | 0.000 |
| C30:1 | 2983 | 0.000 | 0.000 | 0.362 | 0.212 | 0.000 | 0.000 | 0.072 | 0.040 | 0.000 | 0.000 | 0.000 | 0.000 |
| C30 | 3000 | 0.247 | 0.140 | 0.494 | 0.150 | 0.347 | 0.050 | 0.247 | 0.032 | 0.305 | 0.091 | 0.332 | 0.063 |

Table S2: (continued)
| Compound | RI | *A. m. carnica* |  | *A. m. iberiensis* |  | *A. m. ligustica* |  | *A. m. macedonica* |  | *A. m. mellifera* |  | *A. m. ruttneri* |  |
| --- | --- | --- | --- | --- | --- | --- | --- | --- | --- | --- | --- | --- | --- |
|  |  | mean | sd | mean | sd | mean | sd | mean | sd | mean | sd | mean | sd |
| 15-; 14-MeC30 | 3030 | 0.000 | 0.000 | 0.205 | 0.216 | 0.053 | 0.078 | 0.154 | 0.215 | 0.058 | 0.122 | 0.249 | 0.360 |
| C31:2_1 | 3051 | 0.000 | 0.000 | 0.000 | 0.000 | 0.429 | 0.228 | 0.000 | 0.000 | 0.267 | 0.102 | 0.000 | 0.000 |
| C31:2_2 | 3057 | 0.079 | 0.062 | 2.181 | 1.791 | 0.000 | 0.000 | 0.000 | 0.000 | 0.860 | 0.517 | 0.206 | 0.087 |
| 10-; 9-; 8-C31:1 | 3075 | 10.199 | 4.628 | 10.752 | 5.823 | 11.378 | 3.335 | 9.924 | 3.786 | 12.345 | 4.861 | 6.134 | 3.266 |
| C31 | 3100 | 11.501 | 5.855 | 6.727 | 3.540 | 8.826 | 1.386 | 6.147 | 2.135 | 5.313 | 2.579 | 7.316 | 4.155 |
| 15-; 13-MeC31 | 3125 | 1.113 | 1.687 | 2.045 | 2.397 | 1.498 | 1.027 | 2.189 | 2.337 | 0.829 | 1.108 | 3.151 | 3.946 |
| 11, 17-diMeC31 | 3149 | 0.000 | 0.000 | 0.000 | 0.000 | 0.000 | 0.000 | 0.000 | 0.000 | **0.457** | **0.303** | 0.000 | 0.000 |
| 13, 17-diMeC31 | 3150 | 0.000 | 0.000 | 0.000 | 0.000 | 0.000 | 0.000 | **0.736** | **0.349** | 0.000 | 0.000 | 0.000 | 0.000 |
| C32:2_1 | 3150 | 0.000 | 0.000 | **0.542** | **0.467** | 0.000 | 0.000 | 0.000 | 0.000 | 0.000 | 0.000 | 0.000 | 0.000 |
| C32:2_2 | 3158 | 0.000 | 0.000 | 0.000 | 0.000 | 0.210 | 0.271 | 0.000 | 0.000 | 0.119 | 0.206 | 0.090 | 0.133 |
| 10-; 9-C32:1 | 3173 | 0.517 | 0.330 | 0.562 | 0.331 | 1.005 | 0.406 | 1.172 | 0.416 | 0.343 | 0.307 | 0.749 | 0.379 |
| C32 | 3200 | 0.000 | 0.000 | 0.226 | 0.243 | 0.270 | 0.300 | 0.232 | 0.117 | 0.176 | 0.083 | 0.285 | 0.148 |
| C33:2_1 | 3249 | 0.872 | 0.545 | 0.000 | 0.000 | 3.329 | 1.016 | 4.565 | 2.388 | 2.314 | 1.772 | 1.850 | 2.087 |
| C33:2_2 | 3257 | 0.000 | 0.000 | **1.852** | **1.175** | 0.000 | 0.000 | 0.000 | 0.000 | 0.000 | 0.000 | 0.000 | 0.000 |
| C33:2_3 | 3265 | 0.000 | 0.000 | **0.606** | **0.564** | 0.000 | 0.000 | 0.000 | 0.000 | 0.000 | 0.000 | 0.000 | 0.000 |
| 10-C33:1 | 3278 | 22.528 | 11.631 | 6.226 | 2.871 | 23.930 | 8.950 | 26.130 | 13.351 | 6.975 | 3.279 | 23.700 | 13.826 |
| C33 | 3300 | 2.456 | 1.694 | 1.033 | 0.443 | 2.993 | 0.801 | 2.862 | 0.919 | 1.122 | 0.655 | 2.771 | 1.614 |
| 17-; 15-; 13-MeC33 | 3327 | 0.000 | 0.000 | 1.307 | 1.777 | 1.224 | 1.185 | 2.642 | 2.111 | 0.602 | 0.489 | 2.014 | 1.630 |
| 13, 19-diMeC33 | 3349 | 0.000 | 0.000 | 0.000 | 0.000 | 0.000 | 0.000 | 0.000 | 0.000 | **0.343** | **0.158** | 0.000 | 0.000 |

Table S2: (continued)
| Compound | RI | *A. m. carnica* |  | *A. m. iberiensis* |  | *A. m. ligustica* |  | *A. m. macedonica* |  | *A. m. mellifera* |  | *A. m. ruttneri* |  |
| --- | --- | --- | --- | --- | --- | --- | --- | --- | --- | --- | --- | --- | --- |
|  |  | mean | sd | mean | sd | mean | sd | mean | sd | mean | sd | mean | sd |
| C35:2_1 | 3449 | 0.223 | 0.227 | 1.439 | 3.113 | 1.895 | 0.640 | 3.464 | 1.186 | 0.610 | 0.294 | 2.041 | 1.484 |
| 12-; 10-C35:1 | 3470 | 0.614 | 0.443 | 0.449 | 0.231 | 1.921 | 0.616 | 3.770 | 1.477 | 0.495 | 0.149 | 2.129 | 1.642 |
| C35 | 3500 | 0.000 | 0.000 | 0.000 | 0.000 | 0.000 | 0.000 | **0.406** | **0.218** | 0.000 | 0.000 | 0.000 | 0.000 |
| 19-; 17-; 15-MeC37 | 3722 | 0.000 | 0.000 | 0.000 | 0.000 | 0.000 | 0.000 | **0.193** | **0.153** | 0.000 | 0.000 | 0.000 | 0.000 |

**Table S3:** One-way permutational test of multivariate homogeneity of group dispersions contrasting the variance in the CHC profiles between task performance groups. The permutations (n=1000) were restricted to the Subspecies. Df -degrees of freedom; SS - sum of squares; MS - mean of squares; F - F-value; SES - standard effect size.

|  | Df | SS | MS | F | SES | p-value |
| --- | --- | --- | --- | --- | --- | --- |
| Nurses - Foragers | 1 | 0.15 | 0.150 | 18.291 | 14.983 | 0.001 |
| Residual | 118 | 0.97 | 0.008 |  |  |  |
| Total | 119 | 1.12 | 0.159 |  |  |  |

**Figure S1:**
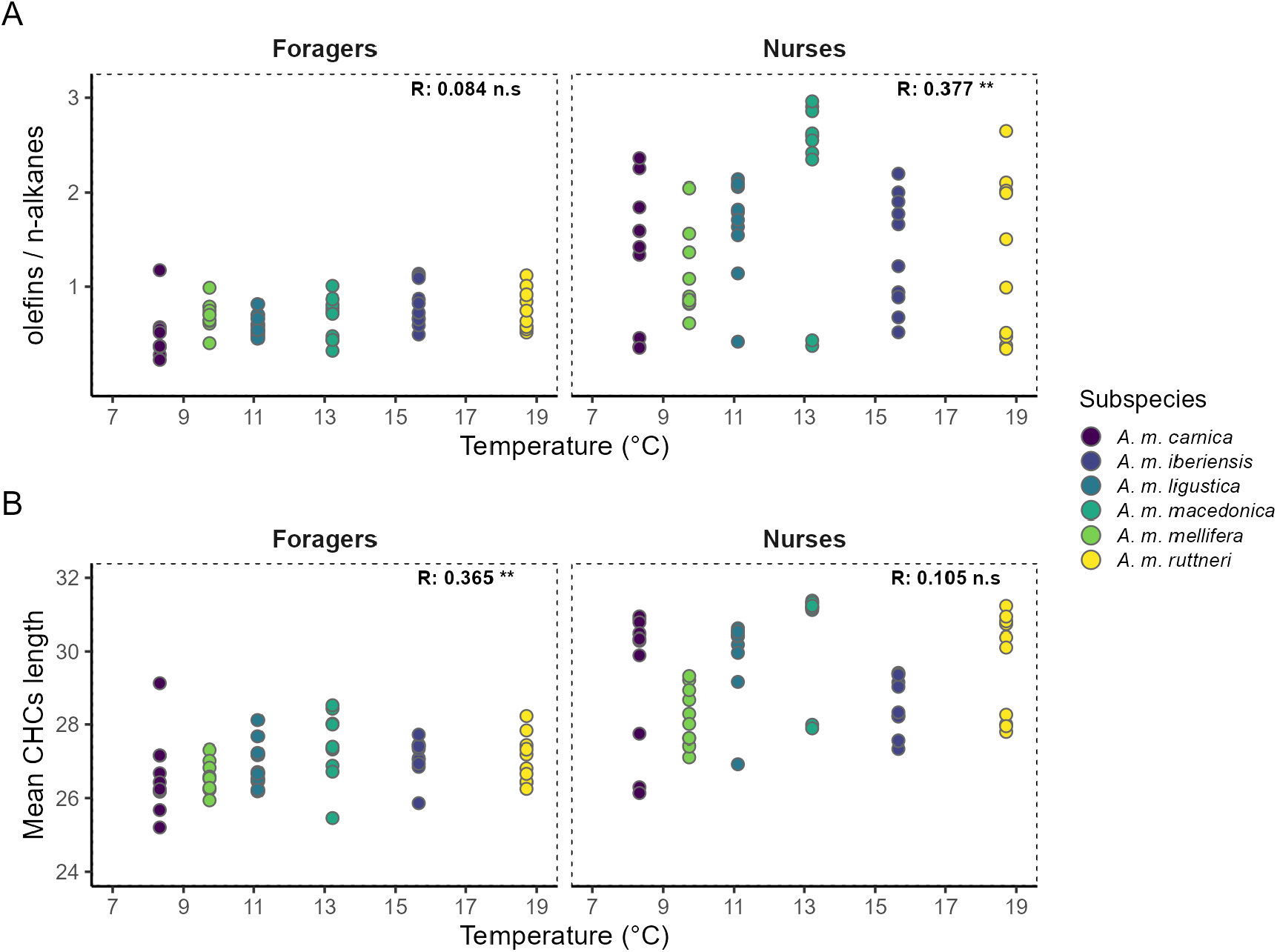
CHC composition vs temperature. A) Spearman’s correlation (R) between the olefins to n-alkanes ratio in the CHC profile of worker bees of different *A. mellifera* subspecies and the average temperature (°C) of the country of origin of the honey bee queens. B) Spearman’s correlation (R) between the weighted mean chain length of hydrocarbons in the CHC profile of worker bees of different *A. mellifera* subspecies and the average temperature (°C) of the country of origin of the honey bee queens. Significance of the correlation is indicated as n.s (p-value 0.05), * (p-value < 0.05), ** (p-value < 0.01), *** (p-value < 0.001)

**Figure S2:**
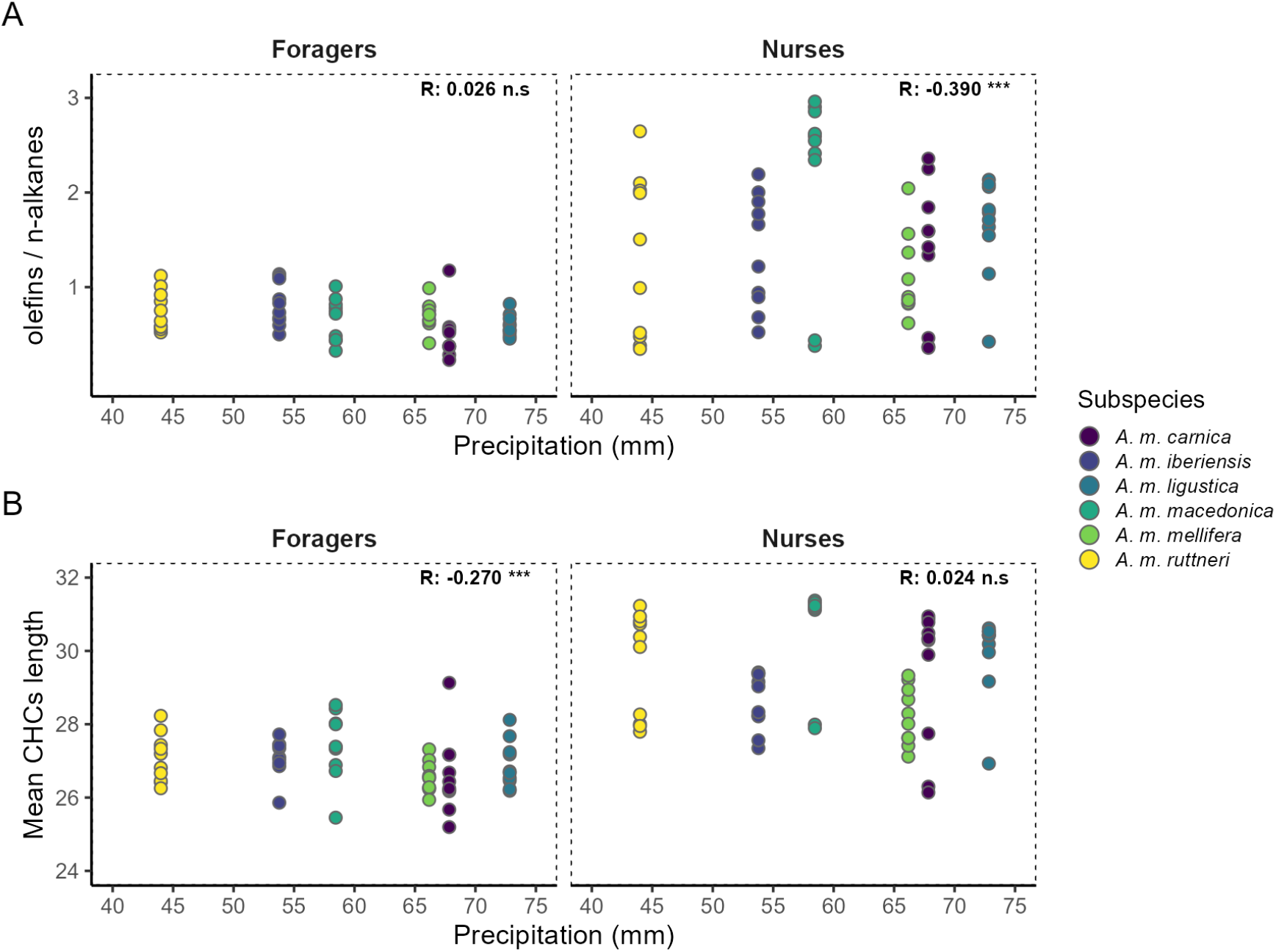
CHC composition vs precipitation A) Spearman’s correlation (R) between the olefins to n-alkanes ratio in the CHC profile of worker bees of different *A. mellifera* subspecies and the average precipitation (mm) of the country of origin of the honey bee queens. B) Spearman’s correlation (R) between the weighted mean chain length of hydrocarbons in the CHC profile of worker bees of different *A. mellifera* subspecies and the average precipitation (mm) of the country of origin of the honey bee queens. Significance of the correlation is indicated as n.s (p-value 0.05), * (p-value < 0.05), ** (p-value < 0.01), *** (p-value < 0.001)

